# Effects of housing conditions and social behavior on methamphetamine self-administration in male and female rats

**DOI:** 10.1101/2025.02.21.639469

**Authors:** Ivette L. Gonzalez, Ammar F. Chauhdri, Reily J. Nessen, Kristin Lee, Jill B. Becker

**Affiliations:** Department of Psychology, University of Michigan; Department of Chemistry, University of Michigan; Michigan Neuroscience Institute, University of Michigan

**Author notes:** One author has been designated as the corresponding author with contact details: Jill B. Becker, Michigan Neuroscience Institute, 109 Zina Pitcher Place, Ann Arbor, MI 48109.

**Keywords:** Social housing, self-administration, sex differences, methamphetamine, substance use, social behavior

## Abstract

Social support is a potentially protective factor against substance use disorders (SUDs). Previous studies in animal models for SUDs have shown that when females are pair housed, they have lower motivation for cocaine and methamphetamine (METH) than females who are single housed. In males, however, social housing has not had the same beneficial effect. This study investigates effects of social housing on METH self-administration in females or males when both cage mates are self-administering METH. The study also investigated how the quality of the relationships changed after METH self-administration. The results show that singly housed females self-administered more METH than socially housed females, while males in both social housing conditions self-administered METH at the same rate. The social behavior data showed that females given saline spent more time apart, however the females given METH spent more time together, suggesting that their social behavior may play a role in the attenuation of METH self-administration. Males’ social behavior remained unchanged after METH and the dominant male in a pair self-administered more METH than the non-dominant male. Females’ self-administration was not affected by dominance. The results of this study show that social housing provides some protective benefits to females, but not males, for METH self-administration. Further, the type of relationship between cage mates affects males’ self-administration and may explain why social housing with a same sex mate is not beneficial for males.

**STRUCTURED ABSTRACT:** 

**Background:** Social support is a potentially protective factor against substance use disorders (SUDs). Previous studies in rats have shown that when females are pair housed, they have lower motivation for drugs such as cocaine and methamphetamine (METH), than females who are single housed. In males, however, socially housed and singly housed males both have the same motivation for drugs such as cocaine.

**Purpose:** The aim of this study was to investigate if social housing would attenuate METH self-administration in females or males when both cage mates are self-administering METH on an intermittent access schedule (IntA). The study also investigated how the quality of those relationships changed after METH self-administration, if dominance played a role in the rate of self-administration, and the extent to which the quality of the relationships was related to METH self-administration.

**Methods:** Male and female rats were individually housed or housed in same sex pairs from around day 42, then animals underwent self-administration of METH (0.3 mg/kg/inf) on an IntA schedule of reinforcement. Both animals in a pair self-administered METH. Social behavior was evaluated prior to and throughout the period when animals were self-administering METH.

**Results:** The results showed that in females, single housed females self-administered more METH than pair housed females, while males in both housing conditions self-administered METH at the same rate. More singly housed females on day one of the IntA had high rates of self-administration, compared to socially housed females. Differences attenuated after the first session, but singly housed females still self-administered METH to a greater extent than pair housed females overall. The social behavior data showed that females given saline spent more time apart while the females given METH spent more time together. This suggested that social behavior may contribute to the attenuation of self-administration in females. Males’ social behavior largely remained unchanged after METH and males continued to exhibit dominance behaviors. Furthermore, males’ METH self-administration was affected by dominance, with dominant males exhibiting greater responding for METH than non-dominant males.

**Conclusions:** Social housing provides some protective benefits to females, but not males, for METH self-administration. Further, the type of relationship between cage mates affects males’ self-administration and may explain why social housing with a same sex mate is not beneficial for males.

**HIGHLIGHTS:**

- Pair housing in female rats resulted in less methamphetamine self-administration compared with individually housed females.
- Pair housing in male rats did not attenuate methamphetamine self-administration compared with single housing.
- Females self-administering methamphetamine spent more time together.
- Males’ self-administration of methamphetamine was affected by dominance status.

## INTRODUCTION

Women escalate from drug use to abuse more rapidly than do men (Becker, 2016; Lynch, 2018). Female rats also exhibit faster acquisition and escalation of drug self-administration, increased motivational withdrawal, and greater drug reinstatement than male rats (Becker and Koob, 2016). Furthermore, females exhibit greater sensitivity to factors contributing to drug intake escalation, such as extended access conditions (Roth et al., 2004; Anker et al., 2009).

Methamphetamine (METH) is a highly addictive stimulant, and the misuse of METH is a continuous problem within the U.S. (Johnston et al., 2015). Females have greater intake and motivation for psychomotor stimulants than males under extended access conditions (Lynch and Taylor, 2005; Corbett et al., 2021). Females’ greater motivation for drugs of abuse, compared to males, remains true in intermittent access (IntA) self-administration paradigms. Females have a more rapid and greater increase in motivation for cocaine than males with IntA (Kawa and Robinson, 2019).

Environmental factors such as social behavior may attenuate self-administration behaviors. For example, social reward consistently prevents drug self-administration across various conditions (Venniro et al., 2022). Studies looking at group housed males compared to individually housed males found that social isolation initially accelerates the acquisition of heroin self-administration, but does not affect cocaine self-administration, suggesting that social isolation is not a necessary condition for the reinforcing effects of these drugs (Bozarth et al., 1989).

While the previous studies focused on social housing and its effect on self-administration, there is little research on whether there are sex differences in the effects of social housing. What research that has been done has shown that pair housing attenuates the escalation of cocaine self-administration motivation in female rats, while isolated males exhibit similar motivation for cocaine as do pair housed males (Westenbroek et al., 2013). When looking at METH self-administration, socially housed females had diminished drug-seeking behavior, compared to single housed females (Westenbroek et al., 2019).

Studies looking at how having a social partner that had self-administered cocaine found that males with an experienced partner were more likely to take drugs compared to if they were exposed to an abstinent partner (Smith et al., 2014). However, not many studies have looked at whether different partners affect females’ drug taking behavior. While we know that there is a sex difference in the effects of social housing in relation to self-administration, in the study reported here we also investigated whether the sex difference remained if both animals were self-administering METH.

How social housing plays a role in drug self-administration is not well understood. One potential factor could be the quality of the relationships between cage-mates. Studies have shown that if males have had an aggressive confrontation with another male, they increase the amount of drug they self-administer (Miczek et al., 1994, 2004, 2008; Ahmed et al., 2020). In this study we investigate the social behaviors engaged in by males and females in the home cage, to begin to understand why there are sex differences in the effects of social housing on drug taking behavior.

## MATERIALS AND METHODS

28 male and 28 female Sprague-Dawley rats (age 40-45 days; arrived from Charles River, Portage, MI) were either housed individually or separated into pairs. Ten females and ten males were housed individually. Nine pairs of females (N=18 females) and nine pairs of males (N=18 males) were matched according to weight and sex on arrival. All animals were placed on a reversed 14h:10h light:dark cycle in a climate controlled colony room. All testing was conducted during the 10 hour dark phase cycle. Animals were given 1 week to acclimate to the colony room before testing began. Food and water were available ad libitum. All experiments and procedures adhered to the National Institutes of Health (NIH) guidelines for laboratory animal use and care, and followed a protocol approved by the University Committee on Use and Care of Animals.

### SOCIAL BEHAVIOR

Two weeks of social behavior were recorded to determine the baseline social behaviors of the rats prior to surgery or METH self-administration (Figure 1). Recordings in this period were conducted daily at the beginning of the dark phase of the cycle for 10 minutes when animals are most active. Animals were recorded in the colony room, but were placed in a new cage for the duration of the recording. Cage-mates remained paired during the recording process. During the self-administration period, animals were recorded for 10 minutes prior to FR1 and extinction sessions. Animals were also recorded at the end of the first session and last session of IntA.

**Figure 1.**
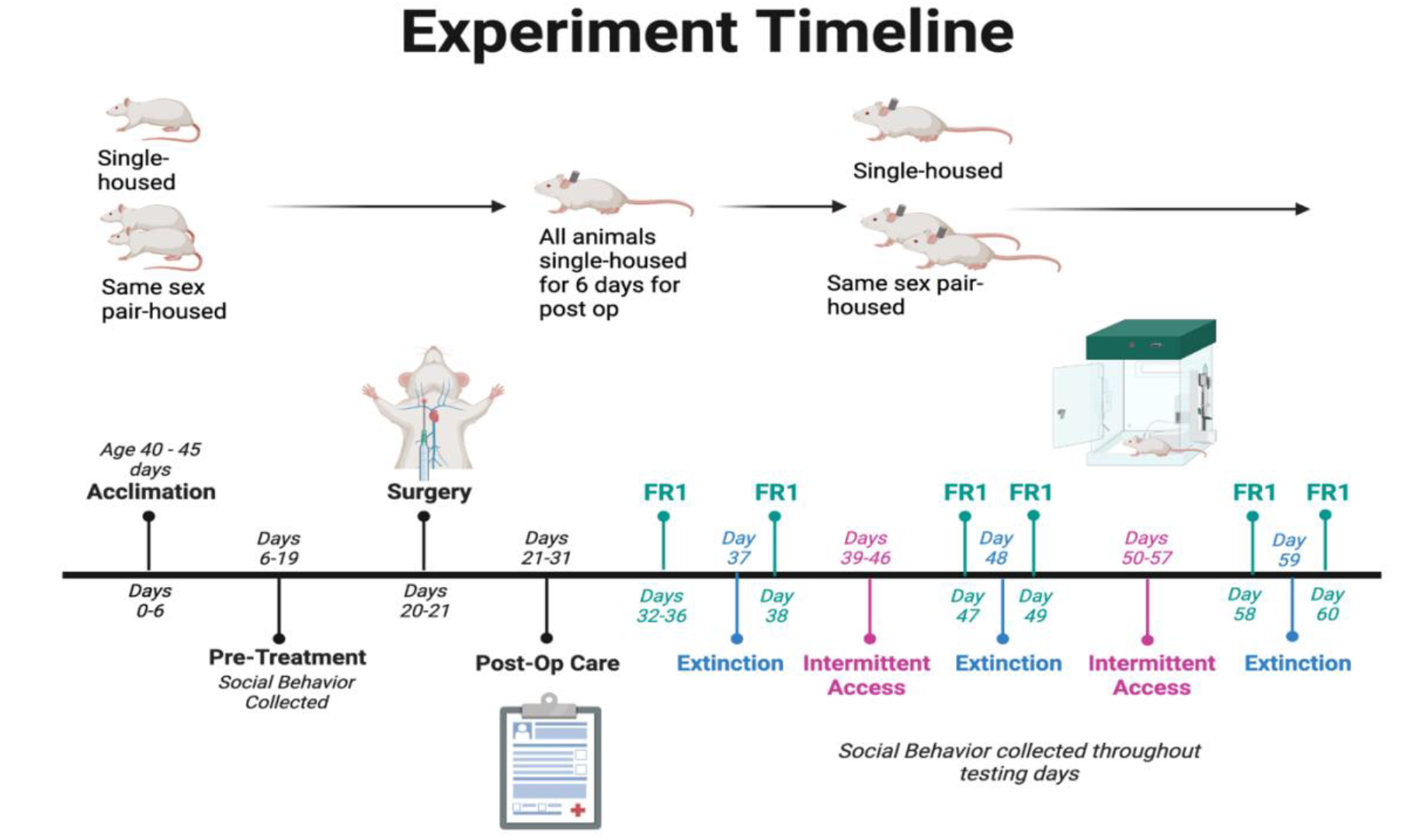
Experimental procedure sequence.

Social behavior videos were coded using BORIS software to label various aspects of rat interactions, including social investigation time, non-social exploration time, and social play time. Additionally, behaviors such as naping, allo-grooming, pinning, and supination were documented for each rat to assess play patterns within each pair. Both the frequency and duration of these behaviors were recorded and analyzed. Results for each behavior and event were compared for each animal pair on the observation recorded days prior to surgery, and the day following each FR1 session for a total of 4 days for each animal.

Dominance was established by looking at the pinning, supine and napping behaviors. Those animals that had higher pinning occurrences were considered the dominant animal and their partner the subordinate. In cases where pinning was very low or the same, naping of the neck was counted and the animal with the higher number of occurrences was named the dominant animal. This was done in order to understand how METH self-administration could be affecting the relationship between cage mates. Data was exported from BORIS into Microsoft Excel, where it was sorted and compiled. Compiled data was then analyzed in GraphPad Prism, where figures were constructed.

### APPARATUS

Self-administration sessions took place in standard operant chambers (Med Associates, Inc.), where animals had the option to nose poke into an active hole for METH or into an inactive hole with no consequences. A cue light was above each nose poke hole, in addition to one house light. Rats were connected to the infusion syringe via a swivel mounted to a counterbalanced arm, allowing them to move freely within the operant chambers.

### CATHETERIZATION SURGICAL PROCEDURE

Following the two weeks of social behavior recording, animals underwent implantation of an indwelling intravenous jugular catheter linked to a backport, a surgery necessary for drug self-administration. These catheters were assembled by adhering silastic tubing to an external guide cannula, and affixing a polypropylene mesh to the base of the cannula to secure the backport in place. The exterior of the backport was shielded by a stainless steel tube surrounding a plastic cap. Rats received a subcutaneous injection of Carprofen (5 mg/kg) 30 minutes before induction of anesthesia with isoflurane (5% isoflurane in oxygen). The distal end of the silastic tubing of the catheter system was inserted into the right jugular vein of the animal and secured using sutures around the tubing and venous tissue. The catheter port exited dorsally from the animal. Following successful implantation, the catheter was flushed with heparin and gentamicin to prevent clotting and infection. Following surgery, animals received Carprofen daily for 3 days (5 mg/kg), and post-operative care was conducted and documented for a total of 10 days. Catheter patency was assessed weekly using an infusion of Brevital (1-5 mg/kg in sterile saline). Surgical procedures and post-operative protocols were reviewed with the appropriate committees prior to surgery.

## SELF-ADMINISTRATION

Animals underwent 5 days of a fixed-ratio one (FR1) training paradigm to learn the audio-visual cues associated with METH availability within the self-administration boxes. After 2 minutes on, the house light would turn off and the lights within the active and inactive nose-poke holes would turn on to signal to the rat the start of the session and drug availability. When the rat poked its nose in the active hole it received a METH infusion (0.06 mg/kg/50 μl inf) over 2.8 seconds, during which time a cue light above the active hole and a tone would turn on. The infusion was followed by a 20-second timeout period during which the cue light and tone would remain on to signal that drug was not yet available. During the infusion and the timeout period the active nose-pokes were recorded but no additional METH infusions were delivered. Following the 20-second timeout period the cue light and tone would turn off to signal that drug was once again available. Throughout the FR1 training, nose-pokes in the inactive port were also recorded but had no consequences. FR1 sessions ended either when the rat reached the maximum of 15 infusions or after 2 hours, whichever came first.

### Extinction (EXT)

Following FR1 training, there was a one-day EXT paradigm which lasted 30 minutes and followed the same procedures as FR1 but no METH was administered. Nose-pokes in the active and inactive ports were recorded. EXT was then followed by another day of FR1 training to reinstate responding.

### Intermittent Access (IntA)

IntA consisted of a 5-minute drug-available period followed by a 20-minute, 30-minute or 50-minute no-drug-available period. The drug-available period consisted of the house light being off and the cue light and tone to accompany METH infusions being received, as in the FR1 training paradigm, though with no timeout period. The no-drug-available period consisted of the house light being on, with nose-pokes being recorded but having no consequences: no cue light, tone, or METH infusions received. Each drug-available period was followed immediately by a no-drug-available period, with each testing day consisting of 8 pairs in succession. No-drug-available period lengths were a randomization of three 20-minute periods, three 30-minute periods, and two 50-minute periods. Before each self-administration session, catheters were flushed with 0.1 ml of sterile saline to verify tubing was not blocked and lavage samples from female rats were collected (male rats were pseudo-lavaged). Following each self-administration session, catheters were flushed with gentamicin (0.2 mL; 3mg/ml) to prevent infection.

### STATISTICS

Statistical analysis was performed using GraphPad Prism. Self-administration data were pooled together by group and examined by session. To analyze the IntA data, a two-way ANOVA was used to investigate the effects of social housing in males and females. A two-way ANOVA was used for extinction data. Groups were compared in 5-minute bins to compare changes in behavior over the testing session. To assess individual differences, rats from each group were assigned to three different categories based on the number of active pokes during drug available and non-drug available periods on the first and last day of each session. Those groups were a rank of low, medium or high (LOW, INT or HIGH). A Kruskal-Wallis test was performed on the ranks of the four groups of animals. A two-way ANOVA was used to determine if there were METH effects on social behavior. A two-way ANOVA was used to analyze how dominance affects motivation for METH self-administration.

## RESULTS: METH SELF-ADMINISTRATION

### FR1

No significant housing or sex differences were found during FR1 within active pokes, inactive pokes and infusions. All animals acquired METH self-administration.

### Effects of social housing on responding during EXT 1

Following acquisition, animals underwent one day of EXT, when active pokes were recorded. Single housed females had higher active pokes compared to socially housed females during the first EXT period ((F (1,31) = 5.484; p = 0.217; Fig. 2; A)). There was also a social housing difference on the first session of EXT in males. Single housed males had higher active pokes compared to socially housed males ((F (1,26) = 5.779; p = 0.0150, Fig. 2; B)).

**Figure 2.**
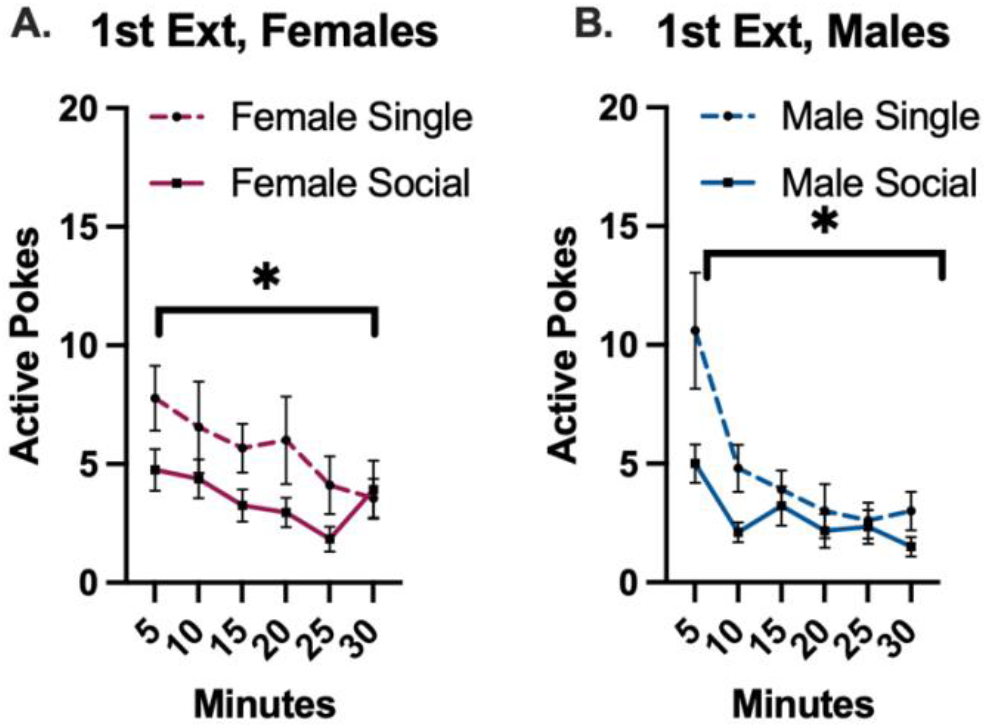
Social housing effects in females and males during the 1st EXT. **A**. Single housed females had a higher number of active pokes in the 1st EXT session compared to socially housed females ((F (1, 31) = 5.848; p = 0.0217)). **B**. Single-housed males had a higher number of active pokes in 1st EXT session compared to socially-housed males ((F (1, 26) = 6.779; p = 0.0150)). *p < 0.05. There were no sex differences.

### Effects of social housing on responding during IA session 1

Responding was reestablished and infusions were recorded during the eight drug available periods during IntA, and totals were compared by each session. Single housed females had a higher rate of infusions during IntA session 1 than socially housed females ((F (1, 26) = 6.948; p = 0.0140; Fig. 3; A)). No social housing effect was found in males (Fig. 3; B).

**Figure 3:**
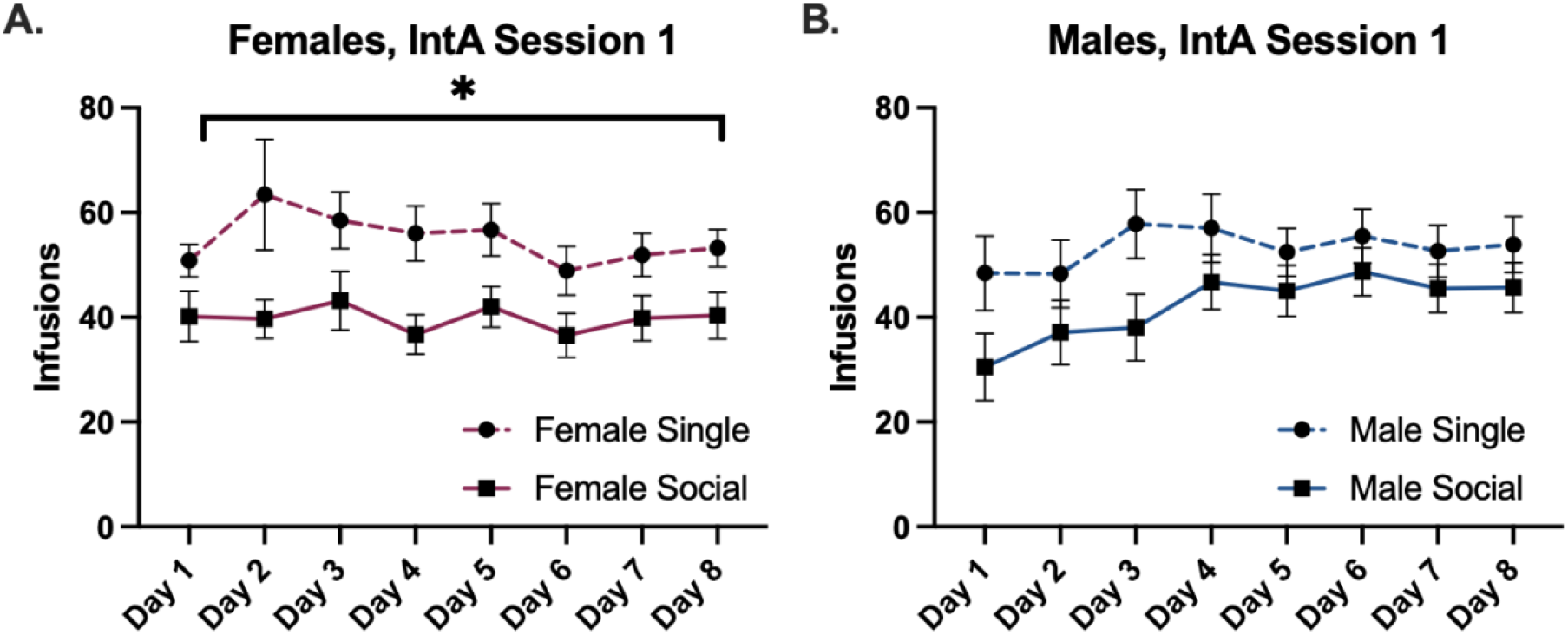
Social housing effects in female and male during IntA session 1. **A**. Single housed females had a higher rate of infusions in IntA session 1. ((F (1, 26) = 6.948; p = 0.0140)). **B**. There was no social housing difference on IntA session 1 in males. *p < 0.05

### Effects of social housing on responding during EXT 2

Following the IntA session 1 was EXT session 2. There were no social housing differences (data not shown).

### Effects of social housing on responding during IA session 2

The animals were then put back on IntA. Single housed females had a higher rate of infusions than socially housed females in IntA session 2 ((F (1, 26) = 6.775; p = 0.0151; Fig. 4; A)). No social housing effect was found in males (Fig. 4; B).

**Figure 4:**
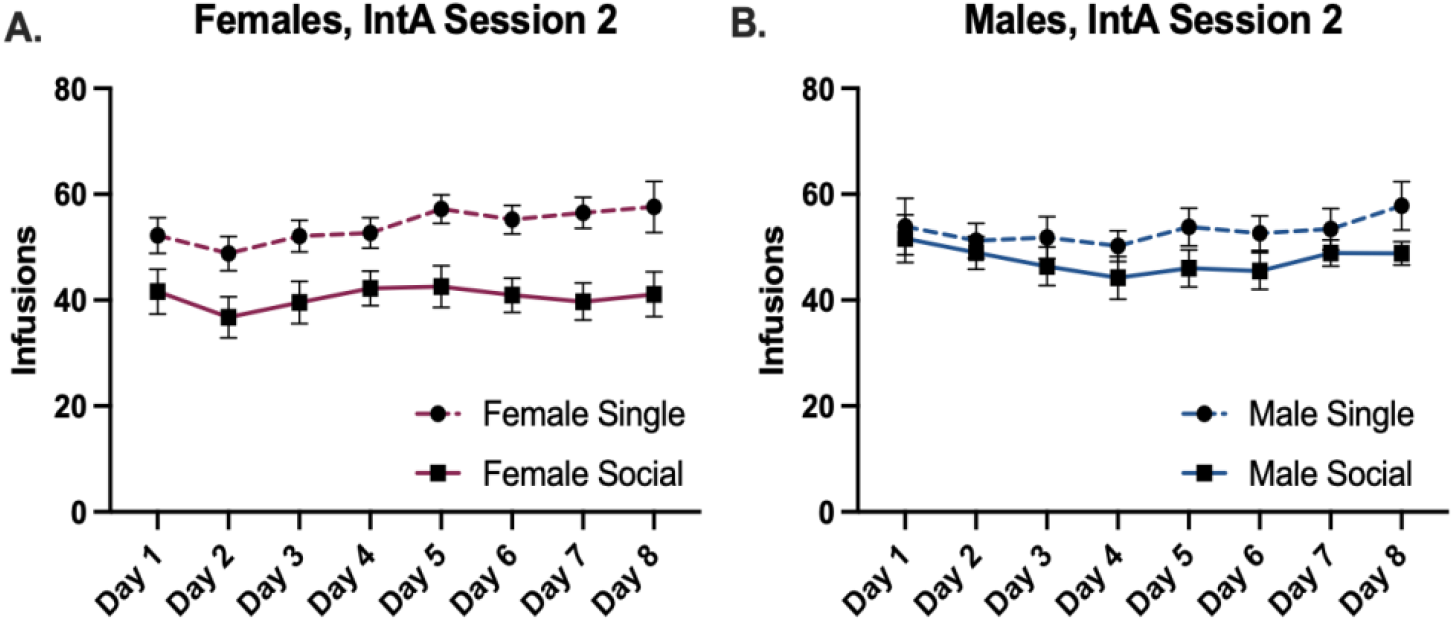
Social housing effects in female and male during IntA session 2.**A**. Single housed females had a higher rate of infusions in IntA session 2. ((F (1, 26) = 6.775; p = 0.0151)). **B**. There was no social housing difference on IntA session 2 in males.

### Effects of social housing on responding during EXT 3

EXT session 3 followed the second IntA period, there were no social housing differences (data not shown).

### Comparison of animal active nose pokes during IntA

The total number of active nose pokes during drug available, and during no drug available periods, was determined for all animals during IntA day one and animals were assigned to low, intermediate (INT) or high (LOW, INT or HIGH) rank. A Kruskal-Wallis test was performed on the ranks of the four groups of animals. The rank totals were significantly different during drug available periods (H (4, n =56) = 10.15; p = 0. 0147; Fig 5; A). On session 1 the number of active pokes during no drug available was higher in single housed females and socially housed males (H (4, n =56) = 9.101; p = 0. 0280, Fig 5; B). In general, social housing increased the proportion of animals in the LOW rank, relative to single housing and this was true for both males and females. There were no significant differences in the rest of the IntA sessions during drug available periods and no drug available.

**Figure 5:**
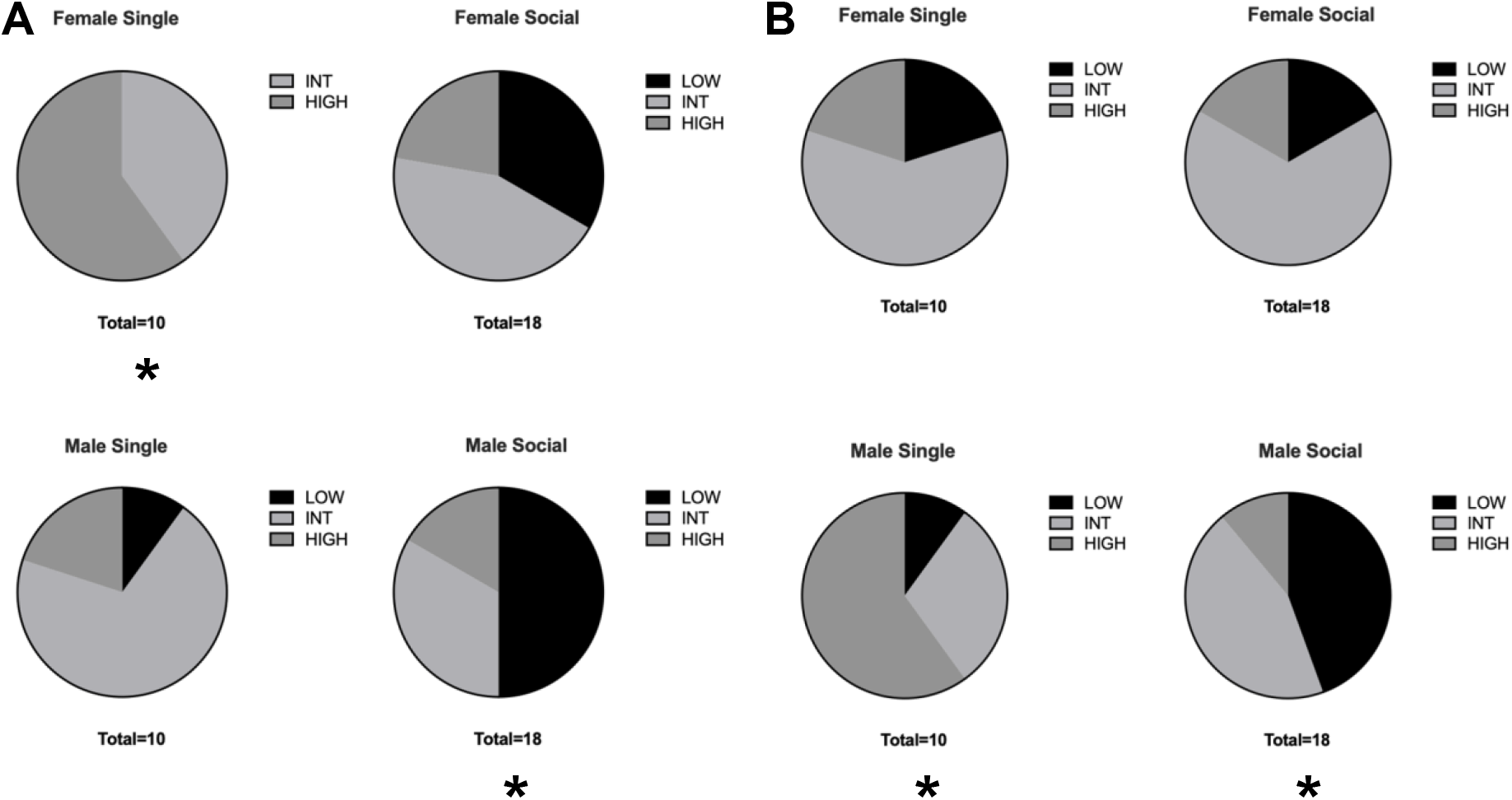
The figures show each social group and the ratio of animals in the LOW, INT or HIGH assignment rank. **A**. Rank of active pokes during drug available on session 1 of IntA. **B**. Rank of active pokes during no drug available period on session 1 of IntA. *p < 0.05

### EFFECTS OF DOMINANCE ON METH SELF-ADMINISTRATION

#### Dominance plays a role in motivation for METH in males but not females

A two-way ANOVA was done to analyze active pokes during drug available periods and no sex differences were found between sessions 1 & 16. There was an effect of dominance in session 1 ((F (2, 42) = 4.566, p = 0.0161); Figure 6). In session 1, dominant males had a significantly higher number of active pokes compared to subordinate males (p = 0.0488; Fisher’s LSD) and single males also had significantly higher active pokes than subordinate males (p = 0.0075; Fisher’s LSD).

**Figure 6:**
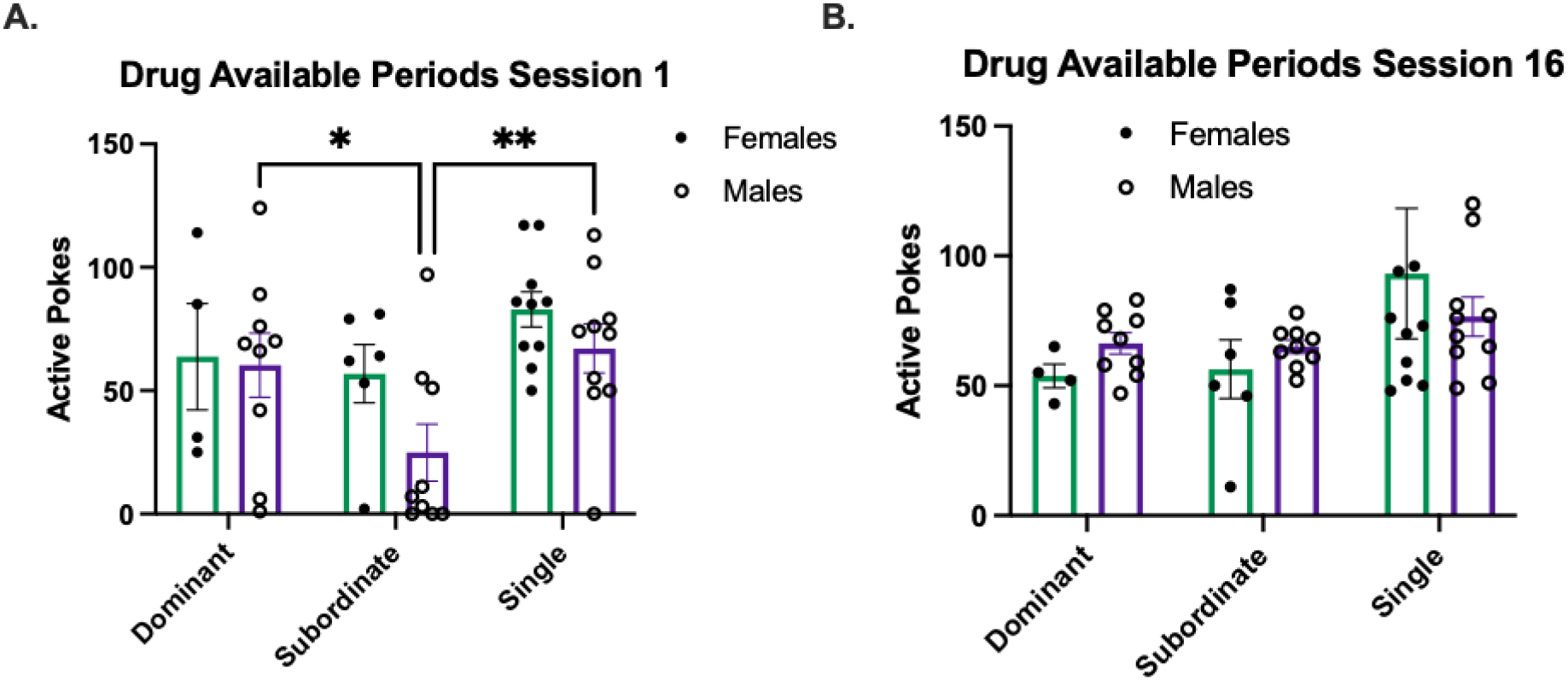
The figures show the number of total active pokes during the drug available periods during session 1 & session 16 of IntA. **A**. No sex differences found, however dominant and single males had a higher number of active pokes compared to subordinate males. **B**. No sex differences or dominance effect found. *p < 0.05, **p < 0.001.

**Figure 7:**
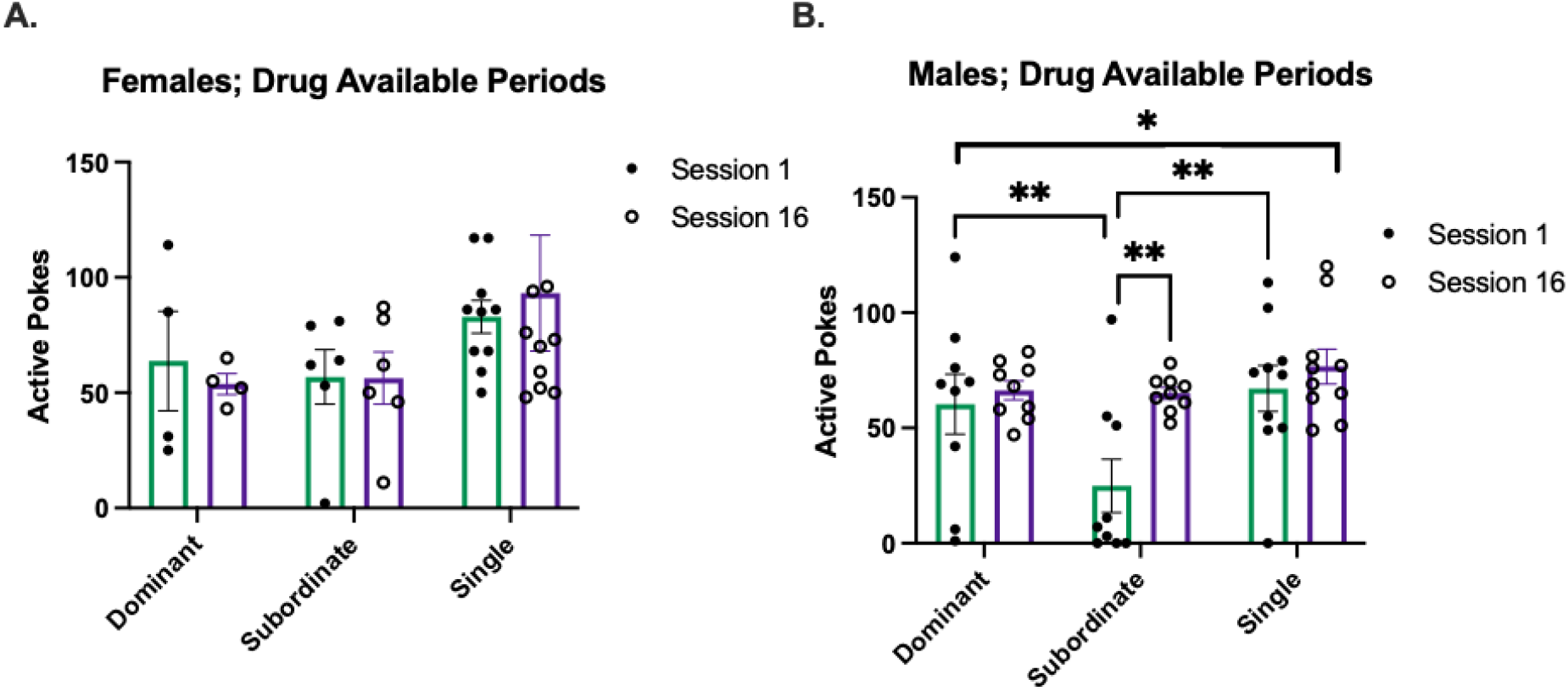
Total active pokes during the drug available periods during session 1 & session 16 of IntA in females and males. **A**. No differences found in females. **B**. In males, there was an effect of dominance and across sessions. Subordinate animals significantly increased total active pokes from session 1 to session 16. In session 1, dominant males and single males had higher active pokes than subordinate males but had no difference by session 16. *p < 0.05, **p < 0.001

We saw similar trends in the no drug available periods. While there were no sex differences, there was an effect of dominance. Using 2-way ANOVA, it shows males in session 1 had an effect of dominance ((F (2, 42) = 3.480, p = 0.0400)) and no significant effects were found in session 16. There was no effects on females in either sessions. Overall, in males, there was an effect of dominance ((F (2, 25) = 4.344, p = 0.0240)). A multiple comparisons test (Fisher’s LSD) showed that dominant males significantly decreased their number of active pokes during no drug available period compared to session 16 (p = 0.0342). This analysis showed that dominance affects motivation for METH in males.

## SOCIAL BEHAVIOR

Social behavior was analyzed in the home cage, as described in the methods, prior to self-administration (baseline) and then before the three extinction tests.

### Time spent apart

A one-way ANOVA was done to analyze differences in time spent apart over time after METH or saline self-administration compared to their baseline. There was a significant effect of treatment in females (F (1, 14) = 8.052, p = 0.0132; Fig. 8; A). Females that received saline spent more time apart than females that received METH. No significant differences were found in males (Fig. 8; B).

**Figure 8:**
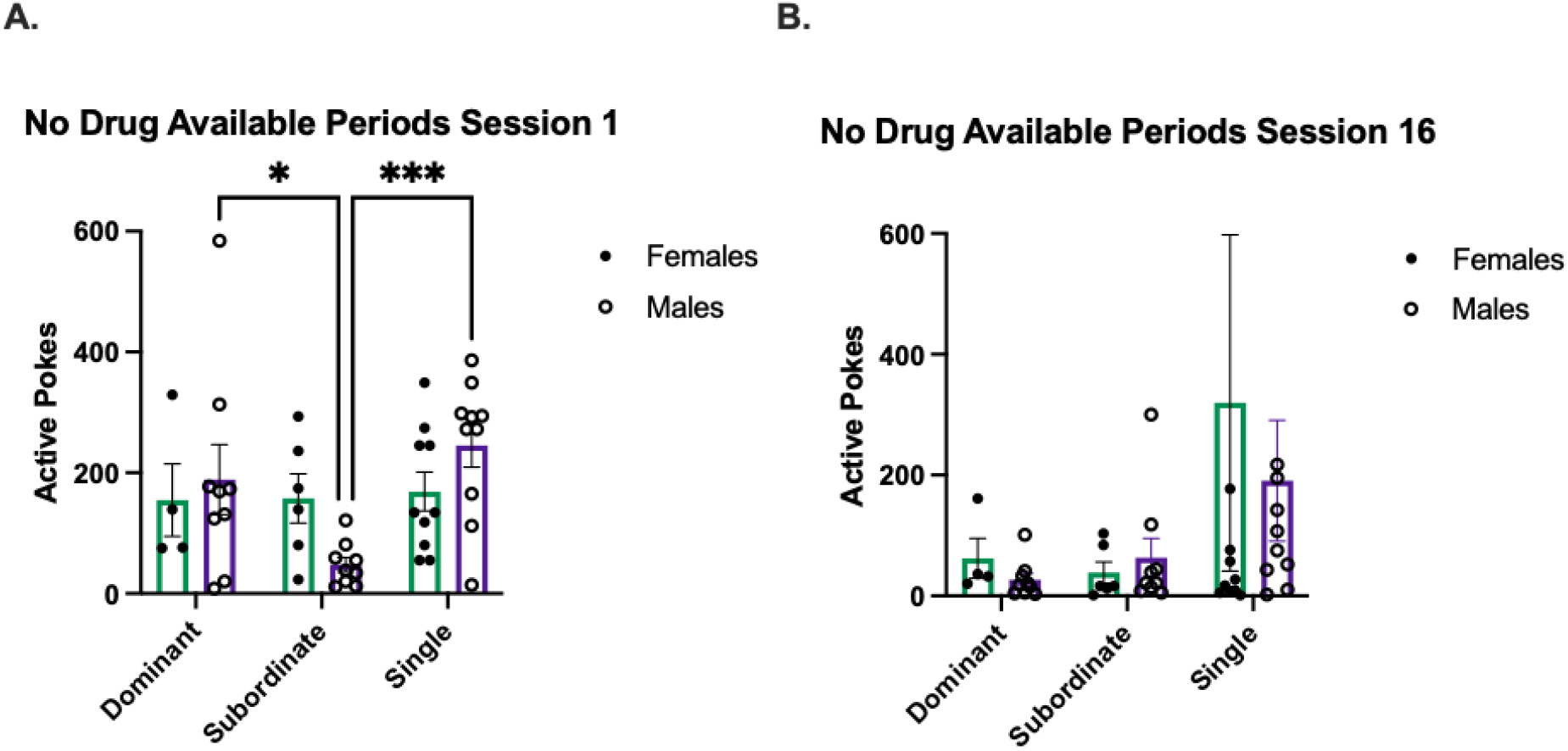
Total active pokes during the no drug available periods during session 1 & session 16 of IntA. **A**. No sex differences found, however dominant males and single males had a higher number of active pokes compared to subordinate males. **B**. No sex differences or dominance effect found. *p < 0.05, ***p < 0.0001

**Figure 9:**
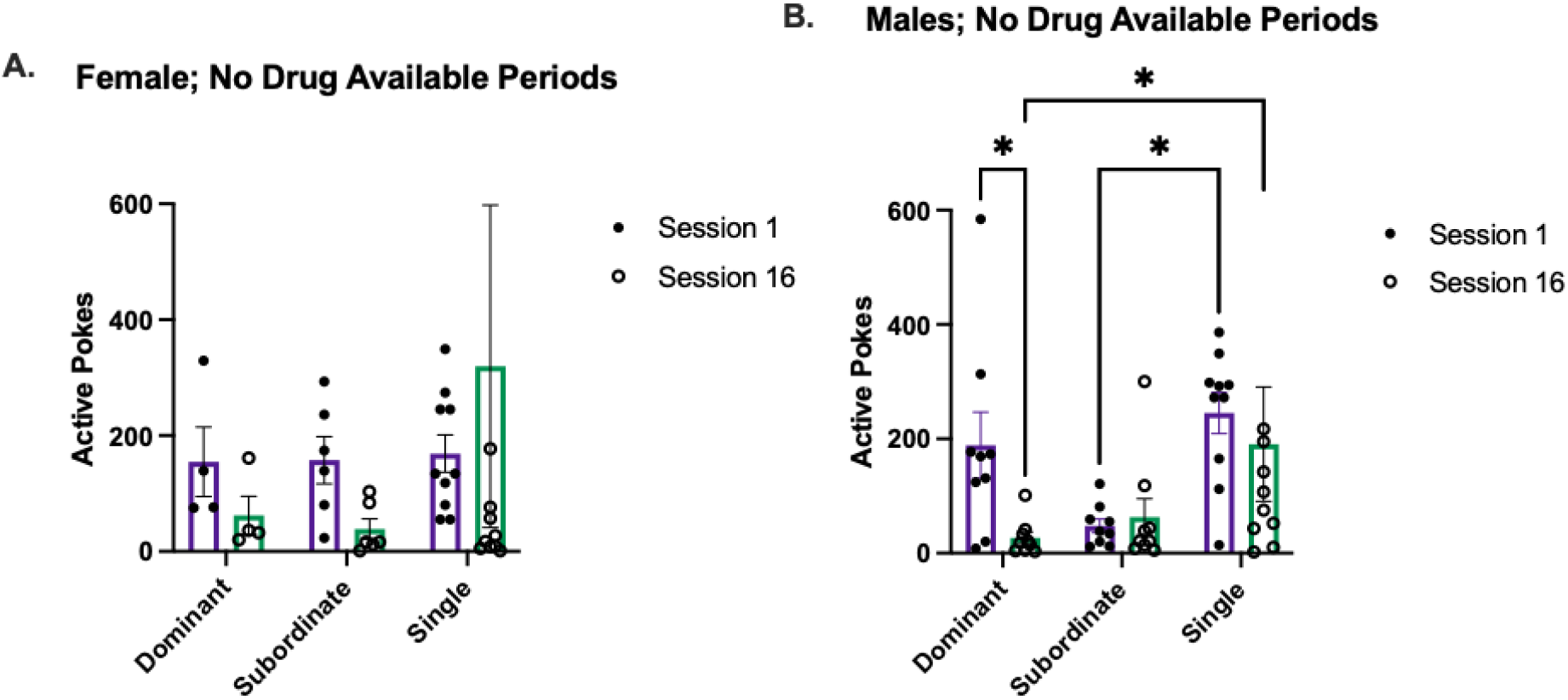
The figures show the number of total active pokes during the no drug available periods during session 1 & session 16 of IntA in females and males. **A**. No differences found in females. **B**. In males, there was an effect of dominance. Dominant animals significantly decreased total active pokes from session 1 to session 16. Fisher’s LSD showed single males had higher active pokes than subordinate males (p = 0.0117) while in session 16, single males had higher active pokes than dominant males (0.0343).*p < 0.05

### Social Interactions

A one-way ANOVA was done to analyze differences in social interactions over time after METH or saline self-administration. No significant difference was found in females (Fig. 10; C). There was a significance of treatment (F (1, 20) = 6.940, p = 0.0159; Fig. 10; D) and time (F (1.911, 38.22) = 4.169, p = 0.0245; Fig. 8; D) in males. Males given saline spent more time interacting with each other than males given METH. Both groups had a decrease in social interactions over time.

**Figure 10:**
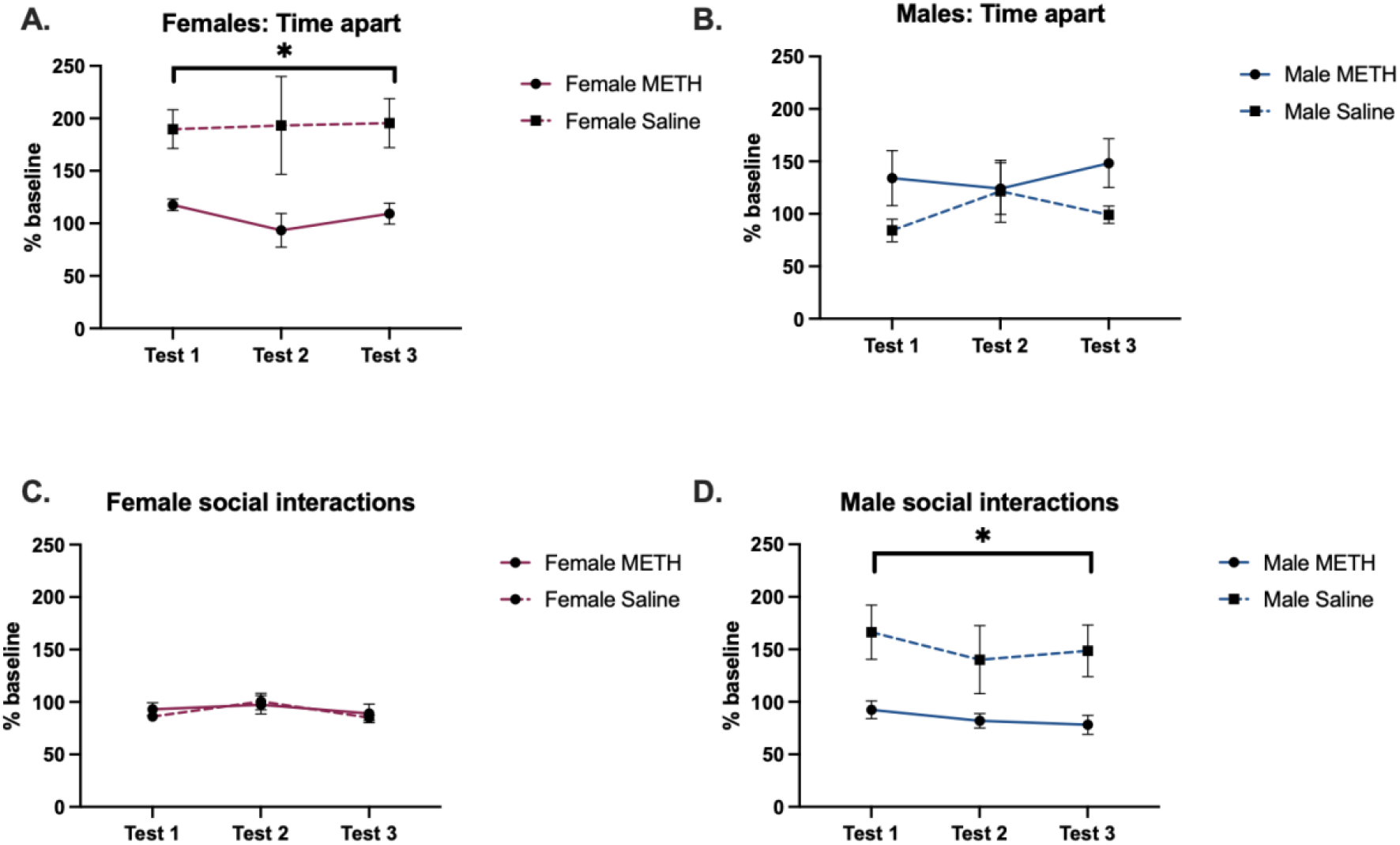
The figures show % baseline for animals that had previously been METH treated or control animals that received saline. **A**. Females spent more time apart across all testing sessions after saline, compared with females that self-administered METH. **B**. No effect of treatment or time was found in time spent apart in males. **C**. No effect of treatment or time was found in females’ social interactions. **D**. Males given saline had more social interactions across all testing sessions than males that self-administered METH. *p < 0.05

**Figure 11:**
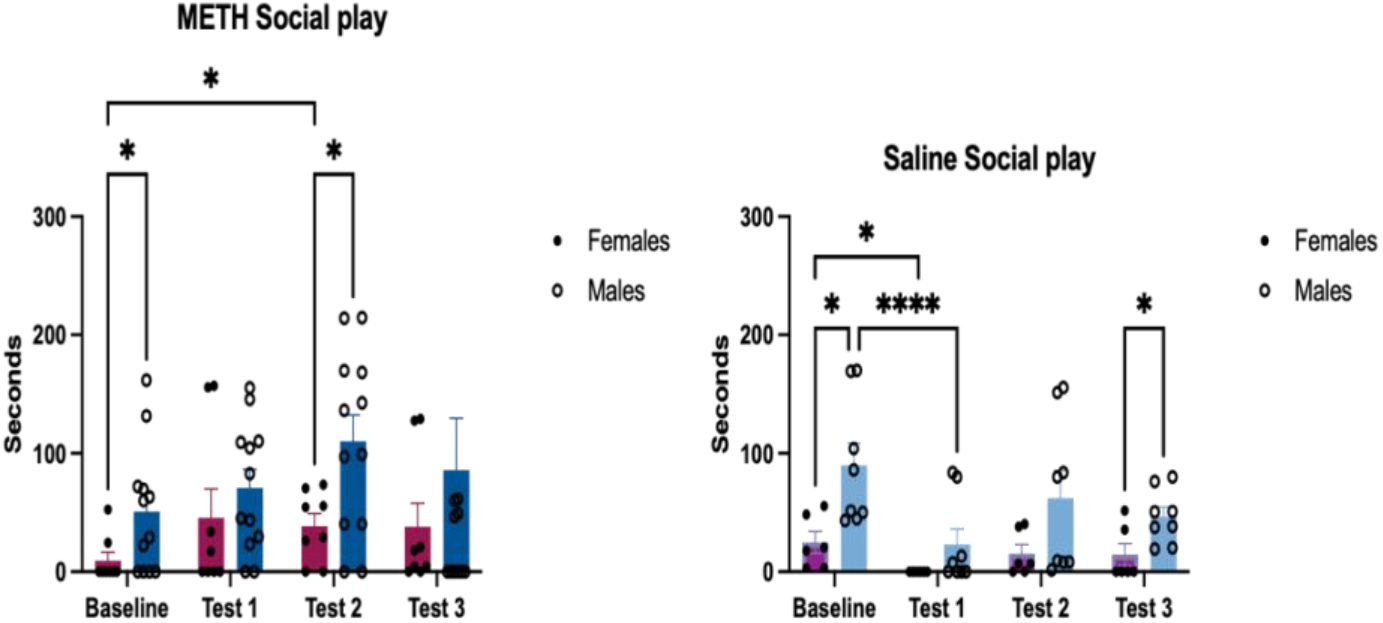
The figures show social play for METH and saline in males and females. **A**. With METH, males spent more time in social play than females at baseline and test 2. **B**. With saline, males spent more time in social play than females and there was an effect of time. *p < 0.05, ****p < 0.0001.

### Social Play

A two-way ANOVA was done to analyze sex differences in social play over time after self-administration. While there were no overall sex differences with animals that self-administered METH, pairwise comparisons showed a sex difference at baseline (p = 0.0279) and at test 2 (p = 0.0103). In animals that received saline, there sex differences in social play (F (1, 12) = 10.35, p = 0.0074) and time (F (1.313, 15.76) = 4.231, p = 0.0477).

## DISCUSSION

Social housing attenuated METH self-administration in females, but there was no effect of social housing in males. This result was true throughout IntA and is consistent with other self-administration studies that have looked at the effects of social housing (Westenbroek et al., 2013, 2019). When looking at individual differences, more single females were in the HIGH category compared to pair housed females, where there were more females in the LOW category, but that difference only occurred on day one. Further studies should continue this paradigm for a longer period of time to assess the long-term effects of social housing in females on METH self-administration.

To better understand why there was a sex difference in the effects of social housing, we looked at social behavior to understand how the relationship between cage mates played a role. Females given METH spent less time apart while males given METH spent less time socially interacting. There was also an effect of social play. Previous studies have shown that social play decreases over time and following the exposure of stimulants such as cocaine (Sutton and Raskin, 1986; Achterberg et al., 2014; Bredewold et al., 2014). Here we see that time does affect social play, as in animals that were given saline social play decreases over time in both males and females. Males overall spent more time doing social play than females when there was no drug introduced. However, after METH self-administration there were no significant sex differences and no difference in time spent doing social play overtime in both males and females. While other studies have shown that social play decreases after cocaine exposure (Achterberg et al., 2014), the results suggest that the psychomotor effects of METH may play a role in not decreasing social play over time. It is possible that the increase in social play can be positive for females but a negative experience for males as pair housing attenuated METH self-administration for females, but not males.

We analyzed the effects of dominance on the motivation for METH and found that it had an effect on males, but not females. During drug available sessions, the subordinate males increased in active pokes compared to dominant males and single housed males. During non-drug available sessions, the dominant and single housed males increased in active pokes compared to subordinate males. Thus, showing that when drugs were available, the subordinate animals were more motivated to poke for drugs but were not when drugs were not available. This is consistent with other findings showing that subordinate monkeys self-administer more cocaine than dominant monkeys (Morgan et al., 2002). Other studies looking at rat cocaine self-administration showed the opposite effect where the dominant rat maintains higher rates of cocaine self-administration compared to their subordinate partners (Jupp et al., 2016), showing that the type of stimulant may have different effects in dominance effect. Further studies need to be conducted to understand the relationship between dominance and self-administration.

## Conclusions

Social support has been shown to be a protective factor for females but not males. Here we found that even when both rats are going through IntA self-administration, socially housed females self-administer less METH than single housed females, while in males there is no effect of social housing. We also saw that dominance played a role in males’ self-administration but not females’. The result of this study has implications for how researchers should consider social housing when doing self-administration studies. The effects found in those studies could be altered depending on social housing. Further studies should look at the effects of group housing compared to pair housing in females and males. Understanding the relationship between cage-mates helps us better understand what component of the social support system is important for reducing drug self-administration. Eventually this work can be used to improve social support systems for humans.

